# Toll-like Receptor 4 Contributes to PCOS-like Metabolic and Reproductive Pathogenesis

**DOI:** 10.1101/2025.10.16.682907

**Authors:** Kiara Wiggins, Zena Del Mundo, Julio Ayala Angulo, Nandini Naidu, Angelina Saba, Christopher Garcia, Christy Nguyen, Naveena Ujagar, Jamie-Jean De La Torre, Gabriela De Robles, Angie Rivera, Davina Trinh, Roberto Tinoco, Alexander S. Kauffman, Varykina G. Thackray, Anshu Agrawal, Marcus Seldin, Gabriela Pacheco Sanchez, Dequina Nicholas

**Affiliations:** Department of Molecular Biology and Biochemistry, School of Biological Sciences, University of California Irvine, Irvine, CA, United States; Department of Biological Chemistry, University of California Irvine, Irvine, CA, United States; Center for Epigenetics and Metabolism, University of California Irvine, Irvine, CA, United States; Department of Microbiology and Immunology, School of Medicine, University of Nevada, Reno, Reno, NV, United States; Department of Obstetrics, Gynecology, and Reproductive Sciences, University of California San Diego, La Jolla, CA, United States; Center for Obstetrics and Gynecology Research Innovation, University of California San Diego, La Jolla, CA, United States; Division of Basic and Clinical Immunology, Department of Medicine, University of California, Irvine, Irvine, CA, United States; Department of Biology, Eberly College of Arts and Science, West Virgina University, Morgantown, WV 26501

## Abstract

Polycystic ovary syndrome (PCOS) is a reproductive disorder with heterogeneous symptoms and severity. Despite extensive research documenting chronic immune dysfunction as a hallmark of PCOS, the specific molecular mechanisms driving immune activation and its connection to the syndrome’s diverse symptoms remain poorly understood. Emerging evidence suggests that gut-derived bacterial endotoxins, particularly lipopolysaccharide (LPS), may breach the intestinal barriers in PCOS patients and trigger systemic inflammation through Toll-like receptor 4 (TLR4), a pattern recognition receptor of the innate immune system. This study investigated whether TLR4 serves as a critical mechanistic driver of PCOS pathogenesis by examining the effect of genetic TLR4 knockout (TLR4^-/-^) in a letrozole (LET)-induced mouse model of PCOS. Our results demonstrate that TLR4 deficiency reduces many PCOS-like symptoms, including elevated luteinizing hormone, anovulation, and metabolic dysfunction. TLR4 knockout also preserved estrous cycling and fertility, improved glucose tolerance, maintained gut barrier integrity, and reduced inflammatory markers in LET-treated females. These findings establish TLR4 as a key mediator orchestrating PCOS’s multi-system pathology, positioning TLR4 as a critical convergence point rather than affecting individual symptoms in isolation. This novel work reveals that TLR4-mediated inflammation drives multiple PCOS pathologies, opening avenues for targeted anti-inflammatory treatments in women with this disorder.

**Significance Statement:** Polycystic ovary syndrome (PCOS) affects up to 15% of reproductive-age women worldwide. This study reveals that TLR4, an innate immune receptor, is key to the pathophysiology of PCOS-like symptoms in female mice. When TLR4 was genetically deleted, mice treated with letrozole to induce PCOS-like symptoms maintained normal weight, glucose regulation, estrous cycling, and fertility. The improvements coincided with preserved gut barrier breakdown and reduced inflammation. These findings identify TLR4 as a key mediator between gut health, immune activation, and PCOS pathophysiology, suggesting that targeting TLR4 could offer new therapeutic approaches for this common but poorly understood syndrome affecting millions of women.

## Introduction

Polycystic Ovary Syndrome (PCOS) is a complex endocrine disorder globally affecting 10-15% of women of reproductive age(*1, 2*). PCOS is characterized by reproductive and metabolic phenotypes including hyperandrogenism, ovulatory dysfunction, and polycystic ovarian morphology. PCOS also elevates the risk of infertility, type 2 diabetes, and cardiovascular disease(*3*–*5*). Despite its widespread occurrence, PCOS mechanisms remain elusive due to its multifaceted etiology. The syndrome is associated with disruptions in the hypothalamic-pituitary-gonadal (HPG) axis, a tightly-coordinated endocrine system regulating reproduction(*6*). HPG axis dysfunction in this disorder often leads to abnormally high levels of luteinizing hormone (LH) and excessive androgen production by the ovaries, which can disrupt ovulation(*7*). Kauffman et al. established a letrozole (LET)-induced hyperandrogenic model of PCOS in C57BL/6J mice(*8*). In this model, chronic treatment with LET, a nonsteroidal aromatase inhibitor, is initiated during peri-puberty and recapitulates most of the known reproductive and metabolic components of the human PCOS phenotype based on the Rotterdam and NIH diagnostic criteria, providing a valuable tool for studying possible PCOS pathophysiology and underlying mechanisms(*8, 9*).

Systemic inflammatory dysregulation has emerged as a consistent finding in PCOS research(*10, 11*). For example, C-reactive protein (CRP), a hepatic acute-phase protein and broad indicator of systemic inflammation, is consistently elevated in PCOS patients across diverse populations(*10, 12, 13*). Yet, despite evidence of the immune system’s involvement in PCOS etiology, the mechanistic role of chronic immune activation remains incompletely understood(*14*–*16*). Recent evidence suggests that LPS, an endotoxin from gram-negative bacteria, mediates immune activation in PCOS(*17*). Clinical studies consistently report elevated LPS levels in serum of PCOS patients, indicating this bacterial endotoxin breaches the gut barrier in PCOS(*18, 19*). In their review of intestinal barrier dysfunction (unrelated to PCOS), Ghosh et al. synthesized findings that increased gut permeability enhances translocation of LPS from the gut into systemic circulation(*20*). These findings may represent a missing mechanistic connection between chronic immune activation and elevated LPS in PCOS.

Toll-like receptor 4 (TLR4) is a pattern recognition receptor of the innate immune system that serves as the primary cellular target for LPS signaling(*21*). Several studies have shown increased expression of TLR4 within the ovaries in animal models of PCOS along with higher serum levels of cytokines associated with TLR4 activation(*22, 23*). Additionally, TLR4 expression is higher in ovarian cumulus cells from PCOS patients than in control women(*24*). LPS binding activates TLR4, promoting the release of pro-inflammatory cytokines and triggering downstream MyD88-dependent NF-κB signaling(*25, 26*). Thus, TLR4 may serve as the critical molecular bridge connecting gut-derived bacterial endotoxins to the systemic immune activity associated with PCOS pathogenesis.

Despite PCOS being associated with endotoxemia, elevated TLR4, and chronic immune dysregulation, whether there is a causal role for TLR4 in PCOS pathophysiology has not yet been determined. In this study, we leveraged the established LET-induced PCOS-like mouse model to determine whether genetic knockout of TLR4 could prevent PCOS phenotypes. We compared C57BL/6J (WT) mice with global TLR4 knockout (TLR4^-/-^) mice and demonstrated that genetic knockout of TLR4 alleviates PCOS-like reproductive and metabolic dysfunction. Our exciting results demonstrate that TLR4 is a key driver of PCOS pathophysiology including elevated LH levels, weight gain, and impaired glucose homeostasis. Importantly, we show that genetic deletion of TLR4 preserves ovulatory function and fertility under conditions that would normally induce PCOS-like infertility.

## Results

### TLR4 Knockout Reduces LET-Induced Weight Gain and Glucose Dysregulation

Many women with PCOS have associated metabolic dysfunction, including obesity and glucose intolerance. Given TLR4 upregulation in hyperandrogenic PCOS rodent models and the known association between inflammation and metabolic outcomes(*27*–*30*), we first tested the necessity of TLR4 for development of metabolic phenotypes in a PCOS-like condition using 4-week-old peri-pubertal C57BL/6J (WT) and TLR4^-/-^ female mice treated chronically with LET or placebo for five weeks (**Figure 1A-B**). This LET mouse model is well-established to recapitulate an overweight PCOS-like condition with metabolic dysfunction. LET-treated TLR4^-/-^ female mice gained significantly less weight over the 5-week treatment period compared to LET-treated female WT mice (**Figure 1C-D**). The reduced weight gain observed in LET-treated TLR4^-/-^ mice was not accompanied by detectable changes in fat or lean mass (**Figure 1E, Supplementary Figure 1A-B**). Next, we measured fasting blood glucose (FBG) and observed no differences regardless of treatment or genotype (**Supplementary Figure 1C**). To further evaluate glucose regulation, we conducted a glucose tolerance test (ipGTT). Compared to LET-treated WT females, LET-treated TLR4^-/-^ mice exhibited more efficient glucose clearance, as evidenced by the decreased area of the curve (AOC) (**Figure 1F-G**). Thus, knockout of TLR4 prevented both the unhealthy weight gain and glucose intolerance in the LET PCOS-like model.

**Fig. 1.**
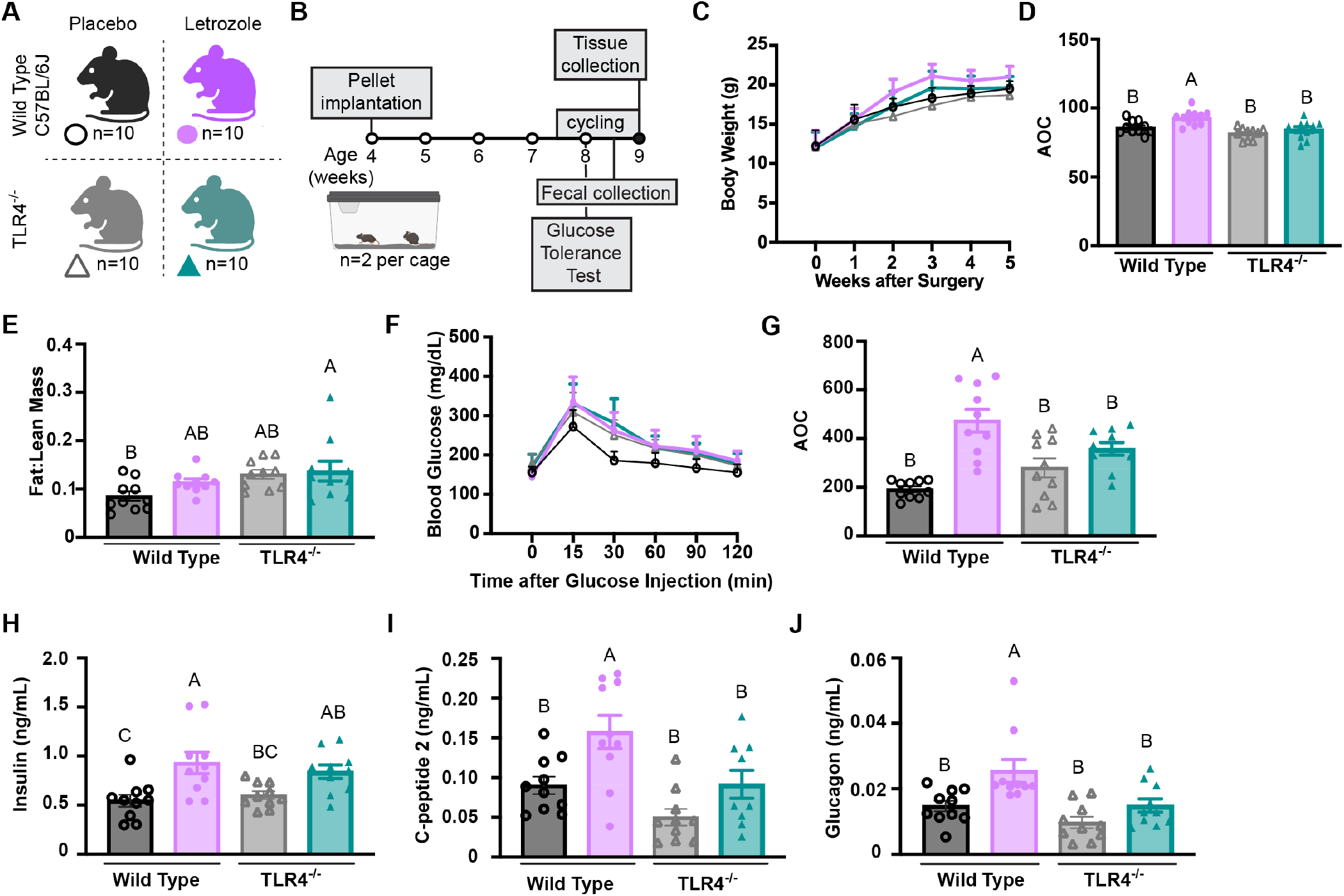
TLR4 Knockout Ameliorates LET-Induced Metabolic and Pancreatic Hormone Dysregulation. (**A**) Overview of experimental groups, colors, and symbols (WT Placebo=10, WT LET=10, TLR4^-/-^ Placebo=10, TLR4^-/-^ LET=10). (B) Timeline of LET or placebo pellet implantation at week 4, with in vivo assessments (glucose tolerance, body composition, estrous cycling) at weeks 8–9 and tissue collection at week 9. (C) Total body weight during the 5 weeks of treatment. (D) Area of the curve for weight gain over a 5-week period. (E) Fat-to-lean mass ratio (% body weight). (F) Glucose tolerance test (GTT) curve and (G) Area of the curve derived from GTT measurements. (H–J) Serum pancreatic hormones: (H) insulin, (I) C-peptide 2, (J) glucagon. Data are presented as mean ± SEM and analyzed with two-way ANOVA followed by Tukey’s post-hoc test. Statistical significance was accepted at p < 0.05 and differences among groups are denoted by a connecting letter system, where groups sharing the same letter are not significantly different from each other, while groups with different letters are significantly different (p < 0.05).

To determine the extent of metabolic protection conferred by TLR4 knockout, we next evaluated fasting serum levels of insulin, its byproduct C-peptide 2, and glucagon to assess pancreatic endocrine function. As anticipated, insulin was significantly higher in LET-treated WT compared to placebo WT mice. In contrast, serum insulin levels trended lower in LET-treated TLR4^-/-^ mice relative to LET-treated WT mice, although this difference did not reach statistical significance. In addition, insulin levels in LET-treated TLR4^-/-^ female mice were not significantly different from their placebo TLR4^-/-^ mice counterparts **(Figure 1H)**. Similarly, C-peptide and glucagon were increased in LET-treated WT mice compared to placebo WT mice, while no increase in these measures was observed in LET treated TLR4^-/-^ mice **(Figure 1I-J)**. To further evaluate metabolic hormone regulation in the absence of TLR4, we measured several gut-derived hormones. Notably, secretin levels were significantly elevated in LET-treated WT mice but were consistently lower in placebo WT mice and remained low upon genetic knockout of TLR4, even upon LET treatment **(Supplementary Figure 1D)**. In contrast, GIP and GLP-1 showed strain-dependent increases in TLR4^-/-^ mice, with LET treatment minimally upregulating serum levels. Resistin levels were not impacted by strain or LET treatment **(Supplementary Figure 1D)**. Collectively, these data suggest that TLR4 deficiency in the context of a LET-induced PCOS-like condition protects from metabolic dysfunction.

### TLR4 knockout prevents PCOS-like reproductive hormone imbalances and estrous cycling disruption

Given that metabolic and reproductive dysfunction are interconnected hallmarks of PCOS(*9*), we hypothesized that TLR4^-/-^ would also prevent the characteristic acyclicity and systemic elevation of androgens and LH observed in the LET PCOS-like model. To test this hypothesis, we evaluated estrous cycles. Supporting prior reports, placebo-treated WT mice maintained normal estrous cycling whereas LET-treated WT mice arrested in diestrus (**Figure 2A**). By contrast, the disruptive effects of LET treatment on ovarian cycling were absent in TLR4^-/-^ mice (**Figure 2A and B**).

**Fig. 2.**
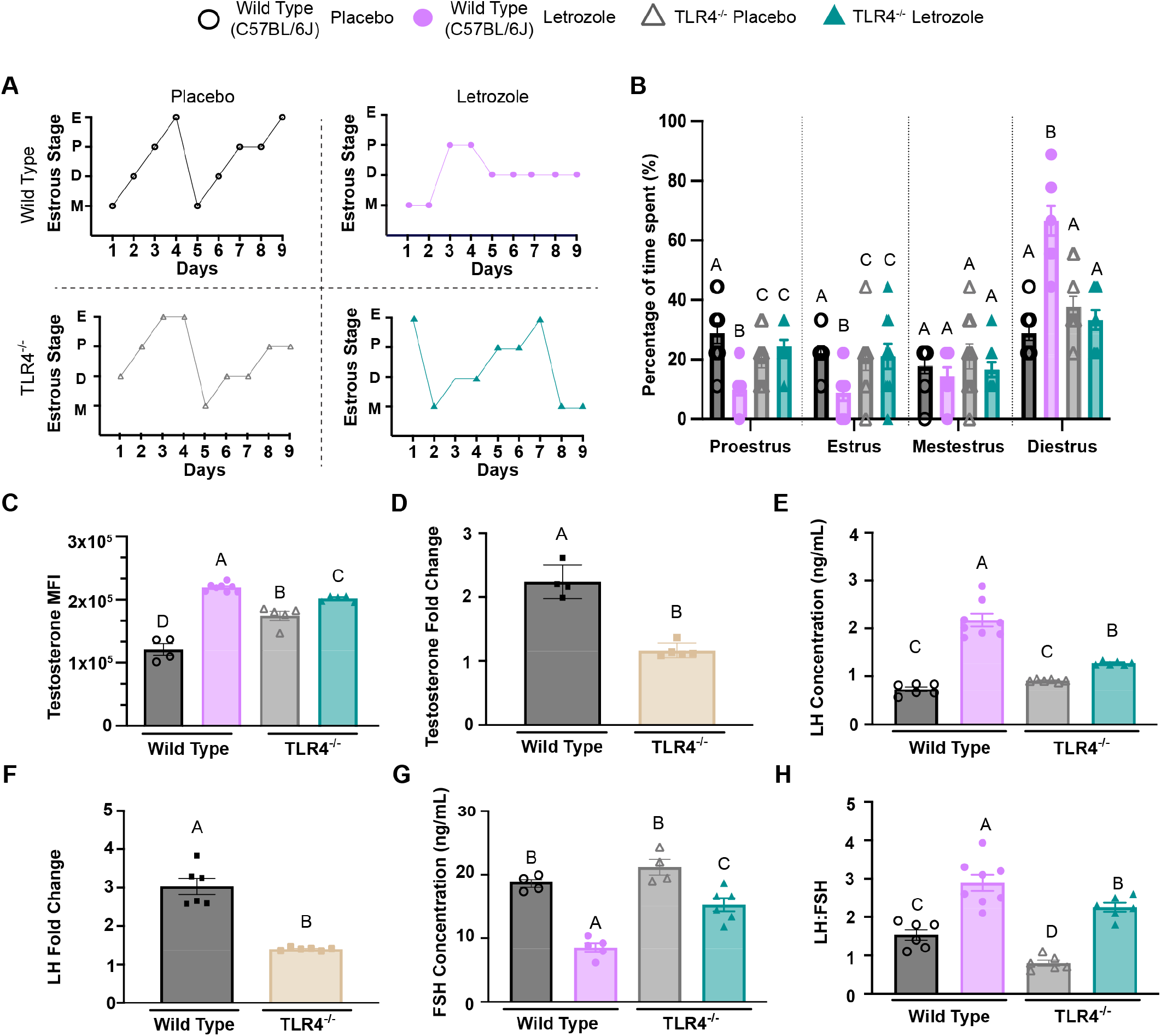
Genetic Knockout of TLR4 Prevents LET-Induced Reproductive Hormone Imbalances and Estrous Cycling Disruption. (**A**) Representative mouse estrous cycling patterns from each experimental group for 9 days. (**B**) Percentage of time spent in each estrous cycle stage (proestrus, estrus, metestrus, diestrus). (**C**) Serum testosterone levels measured by Luminex and displayed as mean fluorescence intensity (MFI). (**D**) Testosterone fold change of WT female mice relative to TLR4^-/-^ female mice. (**E**) Serum luteinizing hormone (LH) concentration (ng/mL). (**F**) LH fold change fold change of WT female mice relative to TLR4^-/-^ female mice. (**G**) Serum follicle-stimulating hormone (FSH) concentration (ng/mL). (**H**) LH:FSH ratio calculated from serum hormone measurements. Data are presented as mean ± SEM and analyzed with two-way ANOVA followed by Tukey’s post-hoc test. Statistical significance was accepted at p < 0.05 and differences among groups are denoted by a connecting letter system, where groups sharing the same letter are not significantly different from each other, while groups with different letters are significantly different (p < 0.05).

To further test the necessity of TLR4 for reproductive PCOS-like phenotypes, we measured serum hormone levels. In WT female mice, LET treatment significantly elevated serum testosterone, as expected based on prior reports(*31*–*33*), mimicking the hyperandrogenemia in PCOS. Testosterone was higher in placebo TLR4^-/-^ mice compared to placebo WT mice. In contrast to the substantial testosterone elevation seen in LET-treated WT mice, LET-treated TLR4^-/-^ mice exhibited only a modest increase (**Figure 2C**). LET treatment increased testosterone more than 2-fold in WT females but LET-induced testosterone change in TLR4^-/-^ mice lower, near1-fold (∼0% increase), owing to the higher baseline (placebo) testosterone levels in this strain(**Figure 2D**). Furthermore, LH levels in LET-treated WTs were higher compared to placebo-treated WT mice (**Figure 2E**), as previously reported(*34, 35*), matching abnormally elevated LH in most PCOS women. By contrast, LET-treated TLR4^-/-^ female mice exhibited significantly lower circulating LH levels compared to LET-treated WT mice (**Figure 2E**), suggesting that TLR4 is necessary for LET-induced increases in LH. LH fold changes remained near 1 in TLR4^-/-^ mice, indicating minimal response to LET treatment, compared to the much larger 2.5-fold increases observed in WT mice (**Figure 2F**). FSH levels were significantly reduced in LET-treated wild-type mice compared to placebo controls, whereas LET-treated TLR4^-/-^ mice showed only a modest decrease relative to their placebo counterparts (**Figure 2G**). The LH:FSH ratio is a common critical indicator of reproductive hormone balance in humans(*9*). LET treatment raised the LH:FSH ratio in WT mice, but the increase was blunted in TLR4-deficient mice (**Figure 2H**). Altogether, these results demonstrate that TLR4 knockout minimizes LET-induced disruption of hormones and reproductive cycling.

### LET-induced PCOS-like ovarian morphology and function is preserved upon TLR4 knockout

To assess whether preserved estrous cyclicity and normal LH levels in LET-treated TLR4^-/-^ mice translated to functional reproductive capacity, we examined ovarian morphology. As expected, LET-treated WT mice showed minimal corpora lutea (CL), indicating impaired ovulation, whereas placebo-treated WT mice and both placebo- and LET-treated TLR4^-/-^ mice displayed multiple CL, indicative of ovulation (**Figure 3A-B**). Furthermore, compared to all other groups, LET-treated WT mice exhibited significantly more cystic follicles (CF), morphological features consistent with PCOS-like ovarian pathology (**Figure 3A-B**). LET-treated WT mice also had higher number of hemorrhagic cysts (HC) than LET-treated TLR4^-/-^ mice (**Supplementary Figure 2B**).

**Fig. 3.**
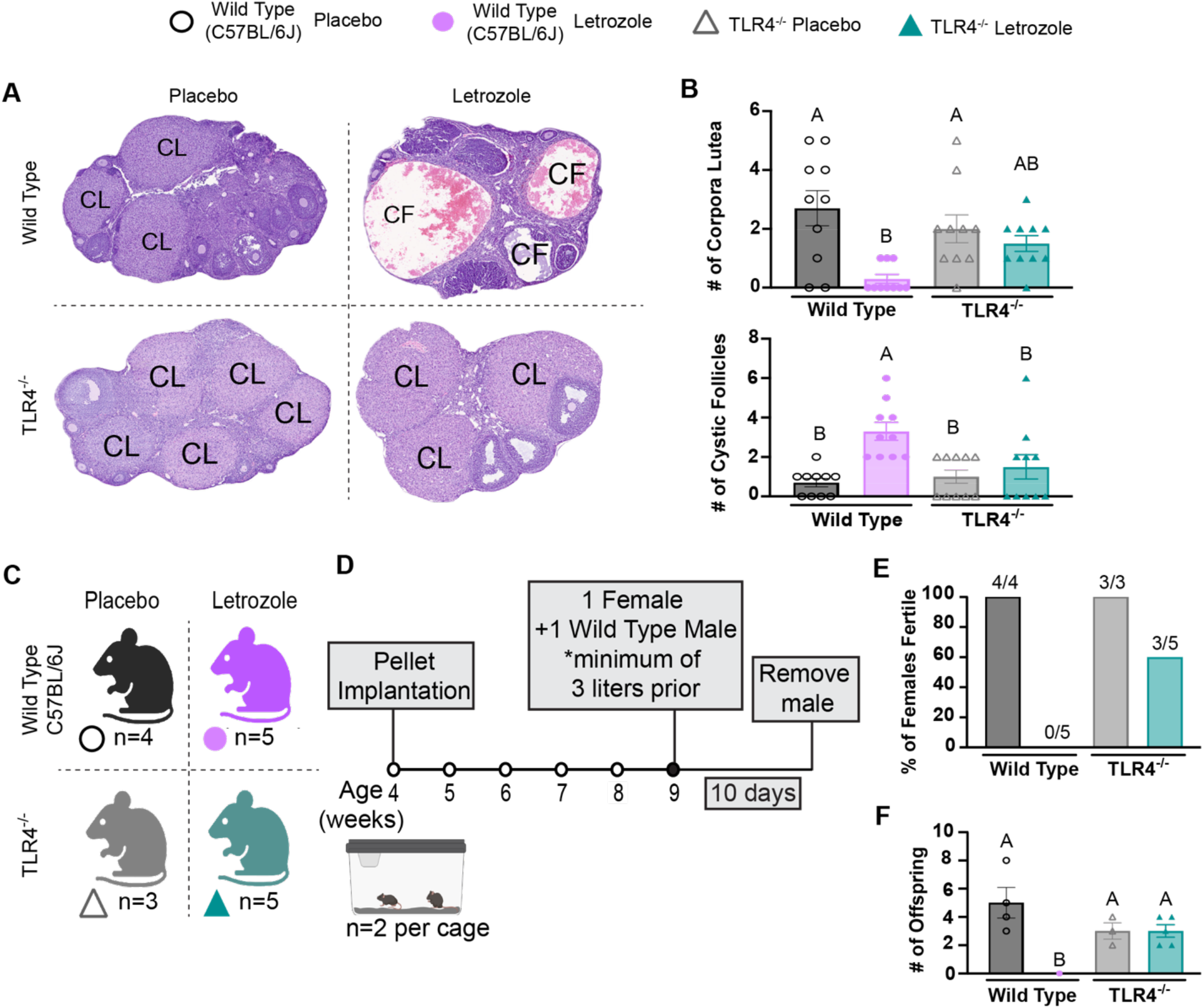
LET-Induced PCOS-Like Ovarian Morphology and Function is Preserved in TLR4 Knockout Mice. (**A**) Representative hematoxylin and eosin (H&E) stained ovarian sections (10 μm) from each experimental group with morphological features labeled: CL = corpus luteum, CF = cystic follicle. (**B**) Quantification of CL and CF per ovarian section. (**C**) Timeline of fertility assessment protocol following treatment period. (**D**) Percentage of fertile females by group (**E**) Average number of offspring per litter from fertile females. Data are presented as mean ± SEM and analyzed with two-way ANOVA followed by Tukey’s post-hoc test. Statistical significance was accepted at p < 0.05 and differences among groups are denoted by a connecting letter system, where groups sharing the same letter are not significantly different from each other, while groups with different letters are significantly different (p < 0.05).

Since the presence of CL in LET-treated TLR4^-/-^ mice is suggestive of preserved ovulation and hence, fertility, we performed a fertility assessment by pairing LET- and placebo-treated WT and TLR4^-/-^ females with proven WT breeder male mice for 10 days, beginning 4 weeks after LET or placebo exposure (**Figure 3C-D**). While placebo-treated WT (4/4) and placebo-treated TLR4^-/-^ female mice (3/3) successfully gave birth to offspring, none of the LET-treated WT females (0/5) were able to produce offspring (**Figure 3E**). Remarkably, 60% (3/5) of LET-treated TLR4^-/-^ mice produced offspring (**Figure 3F**). Though LET-treated TLR4^-/-^ mice produced smaller litter sizes compared to placebo-treated WT mice, we found that the litter sizes were not significantly different between the LET-treated and placebo-treated TLR4^-/-^ mice **(Figure 3F**). These results indicate that TLR4^-/-^ mice showed partial protection from LET-induced infertility, suggesting that TLR4 is necessary for LET-induced PCOS-like reproductive phenotypes.

### TLR4 Promotes Intestinal Barrier Breakdown and Immune Dysfunction

Emerging evidence links gut barrier dysfunction and inflammation to systemic metabolic and reproductive disorders(*36*–*38*). Therefore, we examined if LET treatment induces intestinal inflammation and whether knockout of TLR4 prevents this inflammation. Fecal lipocalin showed modest, genotype-dependent increases in WT mice without statistical significance (**Figure 4A**). In contrast, calprotectin was elevated in LET-treated WT compared to placebo WT mice, while placebo treated TLR4^-/-^ mice had lower levels than WT placebo, with a trending increase upon LET treatment (**Figure 4B**). These findings suggest TLR4 deficiency mitigates against intestinal inflammation. We next investigated whether this local inflammation compromises barrier function using an *in vivo* FITC-dextran permeability assay. LET-treated WT mice exhibited significantly elevated serum fluorescence four hours post-gavage compared to placebo WT mice, while TLR4^-/-^ mice maintained barrier integrity regardless of LET treatment (**Figure 4C**).

**Fig. 4.**
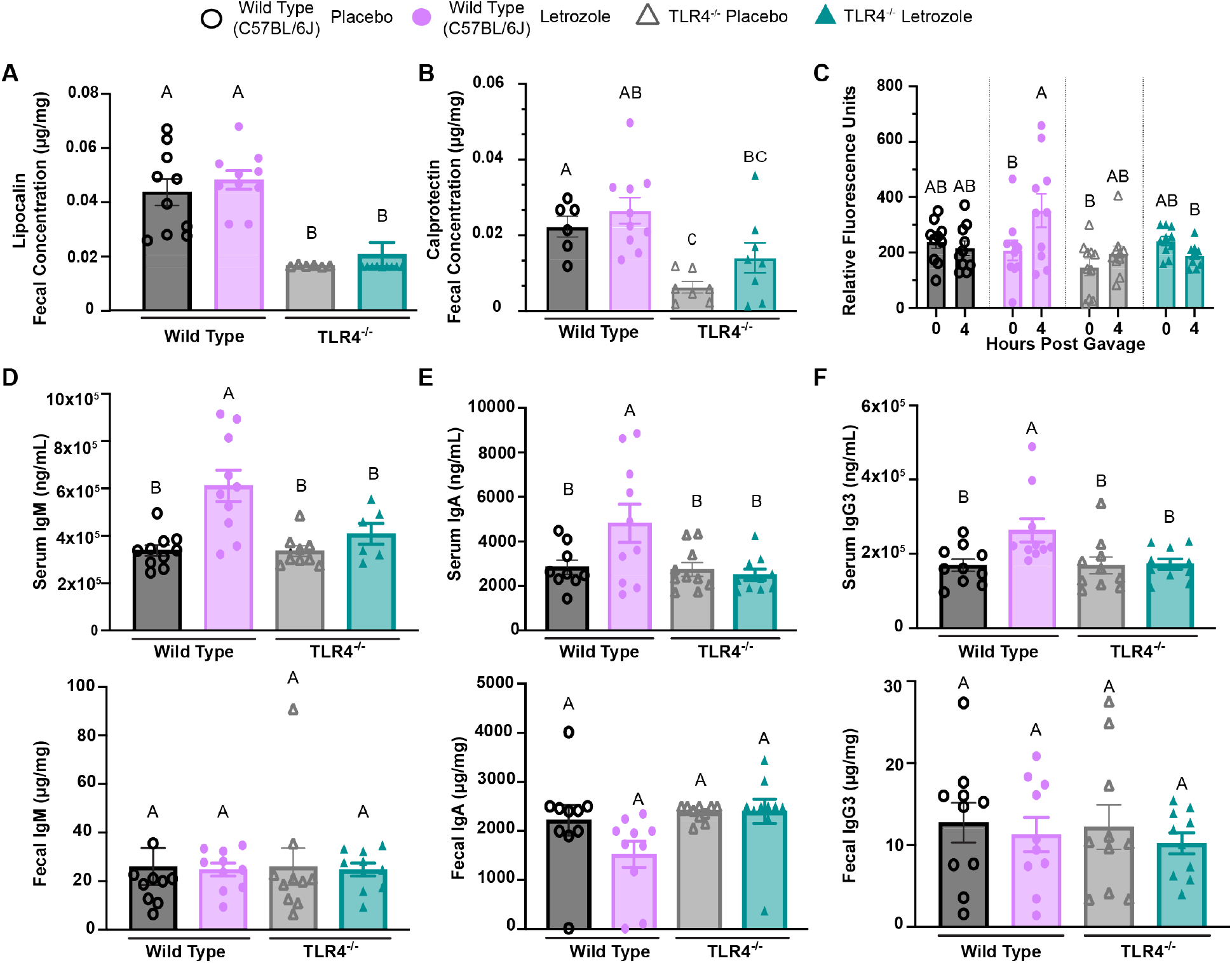
TLR4 Promotes Intestinal Barrier Compromise and Immune Dysfunction. (**A-B**) Fecal lipocalin-2 (A) and calprotectin (B) concentration (μg/mg fecal sample). (**C**) Serum FITC-Dextran fluorescence pre and post 4-hour oral-gavage. (**D**) Immunoglobulin M (IgM) concentration (ng/mL) in serum (top) and fecal supernatant (μg/mg fecal sample) (bottom). (**E**) Immunoglobulin A (IgA) concentration (ng/mL) in serum (top) and fecal supernatant (μg/mg fecal sample) (bottom). (**F**) Immunoglobulin G3 (IgG3) concentration (ng/mL) in serum (top) and fecal supernatant (μg/mg fecal sample) (bottom). Data are presented as mean ± SEM and analyzed with two-way ANOVA followed by Tukey’s post-hoc test. Statistical significance was accepted at p < 0.05 and differences among groups are denoted by a connecting letter system, where groups sharing the same letter are not significantly different from each other, while groups with different letters are significantly different (p < 0.05).

To further characterize immune responses and assess intestinal integrity, we next measured immunoglobulins (Ig) within serum and fecal supernatant. Immunoglobulin M (IgM) was significantly elevated in serum samples of LET-treated WT mice compared to placebo WT mice and TLR4^-/-^ regardless of treatment, indicating overall immune activation (**Figure 4D)**. However, this difference was not observed within fecal supernatant (**Figure 4D**). While IgM serum elevation confirmed systemic immune activation, we next measured IgA to further evaluate mucosal immune activity and barrier function, as IgA is the predominant immunoglobulin at intestinal mucosal surfaces. LET-treated WT mice had elevated serum IgA compared to placebo WT mice and TLR4^-/-^ regardless of treatment, indicating systemic immune activation often associated with compromised barrier integrity (**Figure 4E)**. The reduction of fecal IgA only occurring in LET-treated WT mice is reflective of chronic impaired local mucosal immune function and barrier maintenance(*39*) (**Figure 4E**). Higher concentration of serum IgG3 was only detected in LET-treated WT mice over other groups, but fecal levels trended lower across genotypes based on LET treatment (**Figure 4D**). Together, these results demonstrate that unlike LET-treated WT mice, LET-treated TLR4^-/-^ mice maintained intestinal barrier function and exhibit reduced LET-induced immune activation.

### TLR4 drives shifts in cytokine profiles in LET-induced PCOS mouse model

Given the observed alterations in immunoglobulin levels, we next examined cytokine profiles in both serum and fecal samples to further understand the LET-induced immune response. Heatmaps revealed broad shifts in both serum and fecal cytokine signatures in LET-treated WT mice, which appeared blunted in LET-treated TLR4^-/-^ mice (**Figure 5A and B**). Because inflammatory processes result in the co-variation of cytokines, we further analyzed the cytokine data by partial least squares discriminant analysis (PLS-DA) (**Figure 5C-F)**. For both WT and TLR4-/- mice, serum cytokines could be used to successfully discriminate between placebo and LET treatment with a cross-validation error of ∼68% and ∼82% respectively. Fecal cytokines were unable to produce a discriminatory model, with cross-validation below 30%, indicating low predictive power. The cytokines most important for discriminating between placebo and LET in WT mice as indicated by a VIP score greater than 1 are IL-1β, IL-15, IL-22, IL-23, IL-17F, IL-2, IL-13, and IL-6. Together, these cytokines are characteristic of an innate and adaptive inflammatory response to mucosal injury, such as gut barrier breakdown. When we analyzed the VIP scores in PLS-DA model for TLR4-/- mice, this mucosal injury response was not present. Specifically, IL-5, IL-31, and TNF-α were the major contributors to discriminating between placebo and LET (**Fig 5F**). Interestingly, the LET induced changes in these cytokines are genotype independent (**Fig. 5A-B**).

**Fig. 5.**
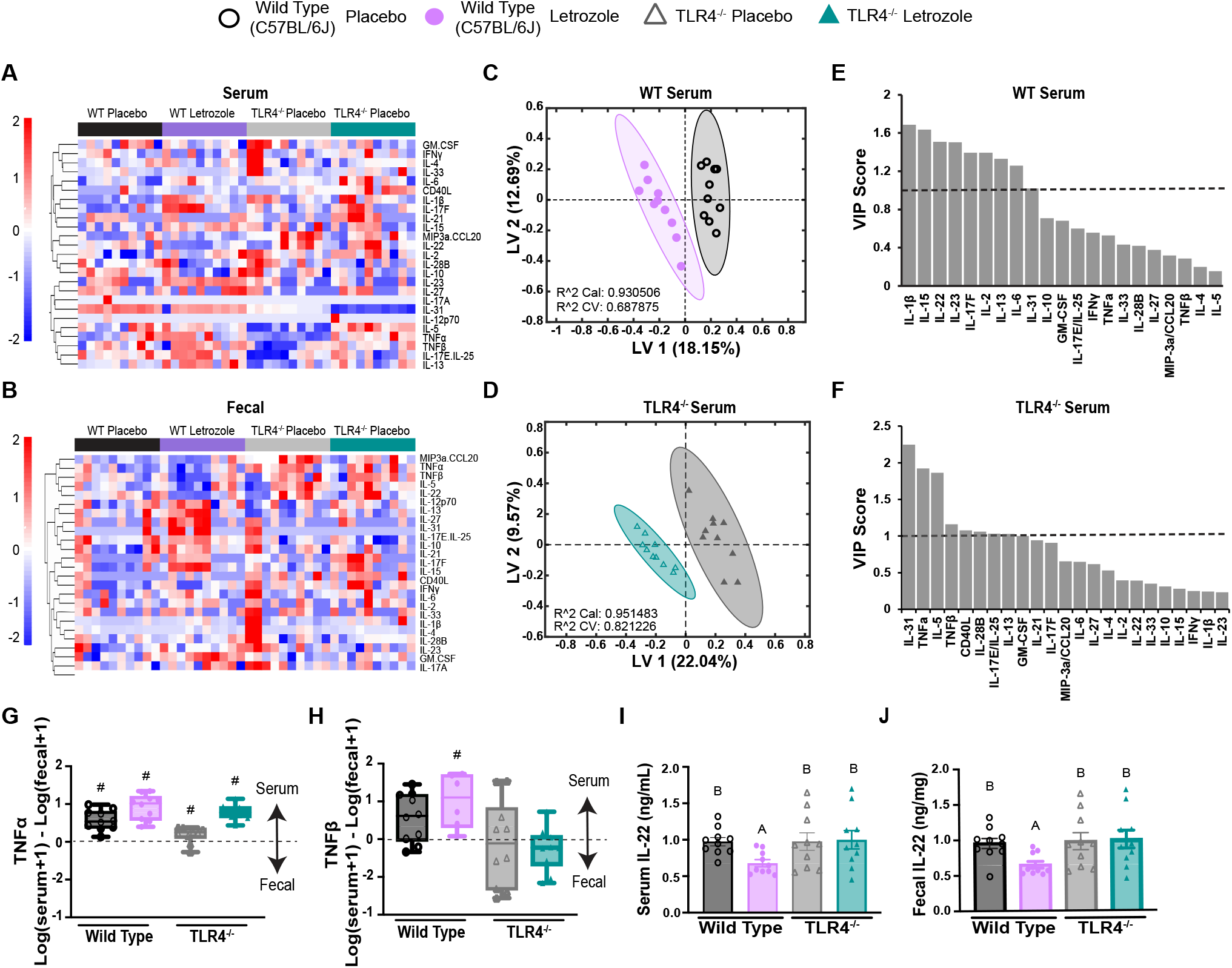
TLR4 Drives Elevated Cytokine Levels in LET-Induced PCOS Mouse Model. (**A-B**) Heatmap of serum cytokine expression profiles from 25-cytokine multiplex panel across experimental groups. (**C-D**) Partial least squares discriminant analysis (PLS-DA) of serum cytokines comparing WT (**C**) and TLR4^-/-^ (**D**) responses to LET treatment. (**E-F**) VIP analysis of cytokine contribution to PLS-DA models in (C and D). (**G-H**) TNF-*α* (G) and TNF-*β* (H) logarithmic ratio of fecal versus serum concentrations (values closer to −1 indicate fecal predominance; values closer to 1 indicate serum predominance). Groups which significantly deviate from 0 are annotated with #. (**I**) IL-22 concentration values in serum (ng/mL) and fecal samples (μg/mg fecal sample). Data are presented as mean ± SEM and analyzed with two-way ANOVA followed by Tukey’s post-hoc test. Statistical significance was accepted at p < 0.05 and differences among groups are denoted by a connecting letter system, where groups sharing the same letter are not significantly different from each other, while groups with different letters are significantly different (p < 0.05).

Given that different cytokine profiles could be used for predictive models for LET treatment in the serum but not fecal supernatant, we evaluated whether individual cytokines preferentially accumulated in the gut versus the systemic circulation. TNF-α significantly favored serum in all experimental groups, suggesting systemic inflammatory responses irrespective of genotype and/or treatment in our model (**Figure 5G**). In contrast, TNF-β only favored serum in LET-treated WT mice indicating that LET treatment in TLR4 competent mice enhances TNF-β-mediated immune responses (**Figure 5H**). Interleukin-22 (IL-22), a cytokine regulated by TNF-β signaling(*40, 41*), decreased in both serum and fecal supernatant of LET-treated WT mice, whereas TLR4^−/−^ mice maintained IL-22 levels under LET treatment (**Figure 5I and J**). Overall, these findings suggest that TLR4 may be an important mediator between gut inflammation and dysregulation of systemic immune responses.

## Discussion

In this study, metabolic, reproductive, and immune dysfunction observed in the LET-induced PCOS-like mouse model was ameliorated upon genetic knockout of TLR4. These findings highlight TLR4 as a central nexus linking the complex phenotypic features of PCOS with intestinal barrier dysfunction and systemic inflammation. As such, TLR4 represents a promising receptor to target in the development of novel therapeutic strategies for this syndrome.

Prior research has established TLR4 contributes to high-fat diet-induced metabolic dysfunction through endogenous lipid activation, adipose tissue inflammation, obesity, and insulin resistance(*42*–*46*). Consistent with this, our findings indicate that TLR4 may have a similar role in hyperandrogenic-driven weight and glucose dysfunction. Interestingly, these improvements were accompanied by alterations in incretin hormones further extending the metabolic benefits of TLR4 deficiency(*47*–*49*). Notably, the metabolic benefits of TLR4 knockout extended to reproductive outcomes, highlighting TLR4 as a causal mediator in PCOS-like reproductive dysfunction beyond prior immune correlations(*14, 22, 50*–*55*). The maintenance of regular estrous cycles in LET-treated TLR4^-/-^ mice indicates that TLR4 signaling can disrupt the coordinated hormonal oscillations essential for reproductive function. Our findings of improved fertility are concordant with previous research showing that TLR4 activation impairs follicular development in ovarian pathophysiology(*56*). Moreover, the protection from elevated LH in LET-treated TLR4^−/−^ mice aligns with evidence that TLR4 activation in theca cells suppresses estrogen production, underscoring TLR4’s role in reproductive hormone dysregulation(*56*). Together, these reproductive and metabolic findings highlight TLR4 as a convergent inflammatory pathway in PCOS-like pathology.

We propose that the metabolic and reproductive improvements observed in TLR4^-/-^ mice result from TLR4-mediated preservation of intestinal barrier integrity, reducing immune dysfunction as highlighted in human studies and PCOS-like animal models(*57*). The elevation of serum FITC dextran fluorescence in LET-treated WT mice provide functional support to previously published data showing tight junction proteins are altered in PCOS-like rodent models, PCOS patients and/or those with obesity(*51, 58, 59*). Intestinal barrier breakdown allows gut-derived toxins such as LPS to enter blood circulation and activate TLR4(*60*). A pilot study in humans found elevation in several parameters of intestinal barrier dysfunction and inflammation in PCOS patients(*61*). Our data support Tremellen and Pearce’s(*62*) PCOS gut barrier-endotoxemia-inflammation mechanism, independent of diet, and *Cani et al’s*(*28*) diet-induced endotoxemia-metabolic dysfunction framework.

The functional changes in intestinal barrier integrity were accompanied by alterations in cytokine signaling that may mediate the TLR4-mediated metabolic and reproductive PCOS-like dysfunction. Specifically, TNF-β exhibited both treatment and genotype specific responses, with TLR4 deficiency reducing the LET-induced elevation of serum-to-fecal TNF-β ratios. Consistent with this, serum IL-22 levels decreased in LET-treated WT mice but were maintained in TLR4^-/-^ mice, aligning with reports of reduced IL-22 in women with PCOS and improvement of metabolic and ovarian function following IL-22 administration in PCOS-like mouse models(*63*). Prior literature shows that lymphotoxin (TNF-β) can interfere with the intestinal epithelial protective functions mediated by the IL-22 pathway(*64*–*66*). Our findings suggest that TLR4 activation may contribute to PCOS pathophysiology by promoting barrier disruption via TNF-β– mediated IL-22 suppression, facilitating antigen translocation and systemic immune activation, consistent with reports in women with PCOS.

The therapeutic potential of targeting TLR4 in PCOS is supported by studies demonstrating that minocycline and emodin, treatments which reduced TLR4 expression, improved metabolic dysfunction, ovarian morphology, and hormonal imbalances(*74, 75*). However, translating TLR4 targets to clinical application will require identification of the specific cell types or tissues mediating these effects, which cannot be achieved with a global TLR4 knockout. Our data strongly implicates gut-mediated mechanisms given the preserved intestinal barrier breakdown and reduced systemic inflammation observed in TLR4^-/-^ mice. Future studies should focus on intestinal-specific TLR4 deletion to establish whether gut-mediated TLR4 signaling is sufficient to drive PCOS-like metabolic and reproductive dysfunction. Overall, our data provides a novel framework for PCOS pathogenesis, implicating TLR4 in the heterogeneity and comorbidities of PCOS and as a potential therapeutic target.

## Materials and Methods

### PCOS-like LET Mouse Model

Twelve Toll-Like Receptor 4 knockout (TLR4^-/-^) mice (RRID: IMSR_JAX:029015) and six C57Bl/6J mice (RRID: IMSR_JAX:000664) were obtained from The Jackson Laboratory and bred in the McGaugh Hall Vivarium at the University of California, Irvine. Mice were maintained under a 12-hour light/dark cycle with ad libitum access to standard chow and water. Two animals were housed per cage, grouped by strain and treatment (DOB: 10/16/2023). At four weeks of age, prior to puberty, female mice were subcutaneously implanted with either placebo pellets or 3-mg LET (LET) pellets (50 µg/day; Innovative Research of America). LET treatment lasted five weeks, following the protocol described(*8*). LET was purchased from Fitzgerald, and custom 60-day continuous-release pellets were manufactured by Innovative Research of America. All procedures were approved by the University of California, Irvine Institutional Animal Care and Use Committee (AUP-21-059).

### Body weight and body composition

Body weights were measured at the beginning of each week, and body composition measurements were taken using the EchoMRI™ Whole Body Composition Analyzer on the 4th week of treatment.

### Estrous Cycle Assessment

The estrous cyclicity was monitored for 9 days prior to euthanasia, approximately 4 weeks after LET or Placebo pellet implantation. Three independent reviewers conducted the assessments, with two reviewers blinded to both the treatment and the genotype of the mice. The stages were classified as follows: Proestrus, characterized by the presence of nucleated epithelial cells; Estrus, by the presence of cornified epithelial cells; Metestrus, by a mixture of cornified epithelial cells, nucleated epithelial cells, and leukocytes; and Diestrus, predominantly by the presence of leukocytes.

### Fertility assessment

A separate cohort of 4-week-old TLR4 Knockout and C57BL/6J (WT) mice was implanted with either LET or Placebo pellets to evaluate fertility. Five weeks after pellet implantation (at 9 weeks of age), LET- or Placebo-treated females (n = 6 per group) were paired with adult C57BL/6J breeder males (average age of 14 weeks, with at least 3 prior successful litters). The presence of vaginal plugs in females was monitored at the beginning and end of their light cycle. After 10 days, the breeder males were removed, and females were observed for litter presence and time to first litter.

### Glucose Measurements

Mice were fasted for 4hr with free access to water. Prior to performing a glucose tolerance test mice were fasted for 4hr with free access to water and weighed for calculation of glucose i.p. injection (Formula: Volume of glucose for injection (μL) = 7.5*body weight (g)). Glucose was prepared as a 20% stock solution. Blood glucose was measured prior to injection and at 15-, 30-, 60-, and 90-minutes post injection. Blood was obtained from a tail tip bleed, and blood glucose levels were measured using glucose strips on a handheld glucometer.

### Serum Collection

Blood samples were collected by tail vein 1- and 3-weeks post pellet implantation. At time of euthanasia, 5-weeks post pellet implantation, blood was collected via cardiac puncture. All blood sample timepoints were allowed to clot at room temperature for 1 h, centrifuged at 2000×g for 10 min, and then serum was collected and stored at −20°C until assayed.

### Fecal Sample Collection and Supernatant Isolation

Fecal samples were collected from mice after 4.5 weeks of LET treatment. Fecal samples were flash frozen in liquid nitrogen immediately after collection. Flash-frozen fecal contents were weighed and reconstituted into a freshly made working solution of 1X phosphate buffered saline and 0.1% Tween 20 at a concentration of 100 mg/mL. This working solution was vigorously pipetted to aid in resuspension. Samples were mixed by vortex at max speed for at 5 minutes until fully homogenized, then centrifuged at 12,000 rpm at 4°C for 10 minutes. The supernatant was transferred to sterile microcentrifuge tubes and stored at −80°C until assayed.

### Fecal Lipocalin and Calprotectin

Lipocalin-2 and Calprotectin levels were measured using R&D Systems’ Mouse Lipocalin-2/NGAL DuoSet ELISA (DY1857-05) and Mouse S100A8/S100A9 Heterodimer DuoSet ELISA (DY8596-05). Absorbance was measured at OD405. All assays were performed according to the manufacturer’s instructions.

### Cytokine Multiplex Immunoassay

A total of 25 cytokines were quantified using the Milliplex MAP mouse Th17 magnetic bead panel (Millipore Sigma MT17MAG47K-PX25) according to the protocol from each respective kit. The analyte concentrations were measured using an Intelliflex instrument (Luminex) and the Belysa software (1.2). Two separate datasets of cytokine quantifications were generated based on whether the analysis was serum or fecal supernatant.

### Tissue Collection

After 5 weeks of LET treatment, mice were weighed, anesthetized with CO2 inhalation, blood collected via cardia puncture, and then decapitated. One ovary from each mouse was fixed in 4% paraformaldehyde at 4°C overnight and then stored in 70% ethanol before histologic processing and the ovary was frozen on liquid nitrogen and store at −80°C.

### Hormone Immunoassays

LH and FSH levels were assessed with the MILLIPLEX MAP Mouse Pituitary Magnetic Bead Panel (Millipore Sigma, MPTMAG-49K). Serum Insulin, Glucagon, C-peptide 2, GIP, GLP-1, Resistin, and Secretin with the Mouse Metabolic Hormone Expanded Panel (Millipore Sigma, MMHE-44K). Serum testosterone was measured via the Multi-Species Hormone kit (MSHMAG-21K) and testosterone flexing pack (SPRCA1825). The Testosterone capture antibody exhibits 100% reactivity to Testosterone and 16% cross-reactivity to 5-alpha-DHT. Briefly, 100 µL of serum was vortexed with 150 µL acetonitrile and incubated for 10 minutes at room temperature. The sample was then vortexed again for 5 seconds, then centrifuged at 17,000 x g for 5 minutes. 200 µL of supernatant was transferred into new Eppendorf tubes. The samples were dried by Speed Vac, followed by reconstitution with 80 µL Luminex Assay Buffer before being analyzed by multiplex assay. Per manufacturer’s instructions, data is analyzed and presented as Mean Fluorescence Intensity (MFI). All assays were performed according to the manufacturer’s instructions.

### Multi-Immunoglobulin Immunoassays

Immunoglobulin levels were assessed with the MILLIPLEX Mouse Immunoglobulin Isotyping Magnetic Bead Panel (Millipore Sigma, MGMMMAG-300K). Assay was performed according to the manufacturer’s instructions.

### Ovarian Histology

Fixed ovaries were trimmed, paraffin embedded, and serial sectioned at 10μm and then stained with hematoxylin and eosin by University of California, Irvine, Experimental Tissue Resource Core. Slides were scanned on a Hamamatsu Nanozoomer scanner. Corpora lutea and Follicular cysts were quantified from the average counts across four serial sections of each ovary from each mouse. Counts were made by an investigator blind to the treatment group.

### In vivo intestinal permeability

FITC Dextran, FD-4 (Sigma-Aldrich, 46944) was prepared in MilliQ (600 mg/kg) and administered via oral gavage in a randomized order. Tail Vein blood was collected pre and 4 hours post FITC Dextran administration. All blood sample timepoints were allowed to clot at room temperature for 1 h, centrifuged at 2000×g for 10 min, and then serum was immediately diluted 1:10 in 1xPBS for a total of 100 µL. Diluted samples were transferred to 96-well black opaque-bottom plate. Relative fluorescence units (RFU) were determined at 530 nm (excitation at 485 nm) using Bio-Rad CFX Duet.

### *Partial least squares* discriminate analysis (PLS-DA)

Partial least-squares discriminant analysis is a supervised analysis approach that uses linear combinations of variables (Placebo or Letrozole) to predict the variation in the dependent variables (cytokines)(*76*–*78*).These analytical tools generate principal components (termed latent variables, or LVs) analogous to those obtained by principal component analysis, but constrained by category (Placebo or Letrozole). Loading analyses ranks dependent variables (cytokines) into LVs that are most important for fit and data separation in the model. Variable importance in projection (VIP) analysis combines all LVs over infinite dimensions. A VIP score >1 is considered important (above average contribution) for model performance and prediction.

All partial least squares analyses were conducted in Solo (Eigenvector Research, Inc.). Data was normalized along each X and Y parameter by Z-score before application of the algorithm. Cross-validation was performed using the leave-one-out strategy. Performance of the feature selection model generated was evaluated by statistics R^2^Cal (calibration) and R^2^CV (cross-validation). A higher R^2^Cal or CV value indicates a better fit or predictive power of the model, respectively. The number of latent variables (LVs) was chosen so as to minimize cumulative error over all predictions. We limited the application of this tool to build a feature selection model using our dataset as a calibration set only, an approach conducted in previously published work(*79*–*81*). Each dataset was z-scored before upload to the Solo software. A model is generally deemed to have significant predictive power to classify with an error rate greater than 70%.

### Serum vs Fecal Cytokine Preference Analysis

Within-mouse comparisons of serum and fecal cytokine concentrations were calculated using log-transformed ratios for each cytokine as log(serum + 1) – log(fecal + 1). The addition of a small constant (+1) prevented undefined logarithmic values from zeros while retaining all observations. Analyses were performed within each experimental group defined by genotype and treatment, evaluating cytokine differences at the individual-mouse level. For each cytokine, a one-sample t test determined whether the mean log difference within a group significantly deviated from zero, where positive values indicated higher serum levels and negative values indicated higher fecal levels. To ensure statistical validity, t tests were conducted only when groups contained ≥5 non-missing paired values and exhibited non-zero variance in log differences. Cytokine-group combinations failing these criteria were excluded from analysis.

### Statistical Analysis

All data were presented as mean ± SEM for each group. Group differences for all datasets were analyzed using Two-Way ANOVA and post-hoc Tukey via JMP Pro 18 software. Graphs were created using R studio 2025.05.1+513 and Graphpad Prism 10. All statistical analyses results with p ≤ 0.05 were considered significant.

## Acknowledgments

We gratefully acknowledge Dr. Nir Drayman from the University of California, Irvine for generously providing the Bio-Rad CFX Duet equipment to measure mouse serum fluorescence. His contribution was instrumental to the success of this research.

## Author Contributions

Conceptualization: KW, DN

Methodology: KW, VGT, DN

Investigation: KW, ZDM, JA, NN, AS, CG, CN, NU, JDLT, GDR, AR, DT

Visualization: KW, ZDM, DN

Supervision: KW, ZDM, DN

Writing—original draft: KW, DN

Writing—review & editing: KW, ZDM, GDR, RT, ASK, VGT, AA, MS, GPS, DN

## Data and materials availability

All data are available in the main text or the supplementary materials

---

**Supplementary Figure 1.**
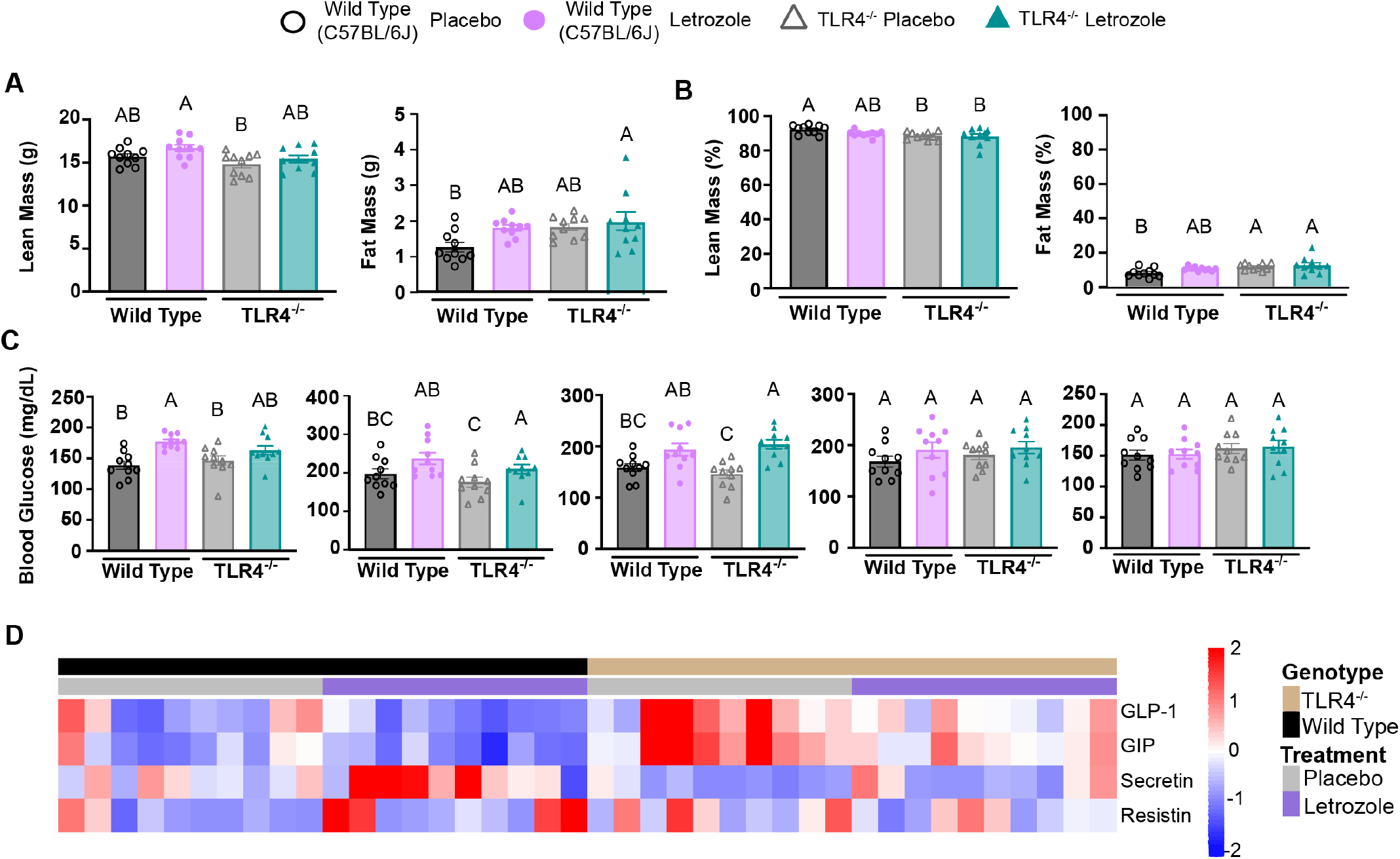
(Related to Figure 1) Body composition and metabolic measurements. (A) Absolute lean and fat mass. (B) Relative lean and fat mass as a percentage of total body weight. (C) Fasting blood glucose levels measured pre-surgery and weekly post-surgery. (D) Heatmap showing relative levels of incretin hormones (GIP, GLP-1, Secretin, and Resistin). The heatmap uses a tricolor scale (red = high, white = intermediate, blue = low expression).GLP-1, group means differed significantly as follows: WT placebo (b), WT LET (b), TLR4^−/−^ placebo (a), TLR4^−/−^ LET (c). GIP: WT placebo (bc), WT LET (c), TLR4^−/−^ placebo (a), TLR4^−/−^ LET (b). Secretin: WT placebo (b), WT LET (a), TLR4^−/−^ placebo (b), TLR4^−/−^ LET (b). Resistin, no significant differences were observed among groups (all a). Group color coding: WT (black) with placebo (gray) and LET (purple); TLR4^−/−^ (tan) with placebo (gray) and LET (purple). Data in panels A–C are presented as mean ± SEM and analyzed by two-way ANOVA with Tukey’s post hoc test. Statistical significance was accepted at *p* < 0.05, with groups sharing the same letter not significantly different from each other.

**Supplementary Figure 2.**
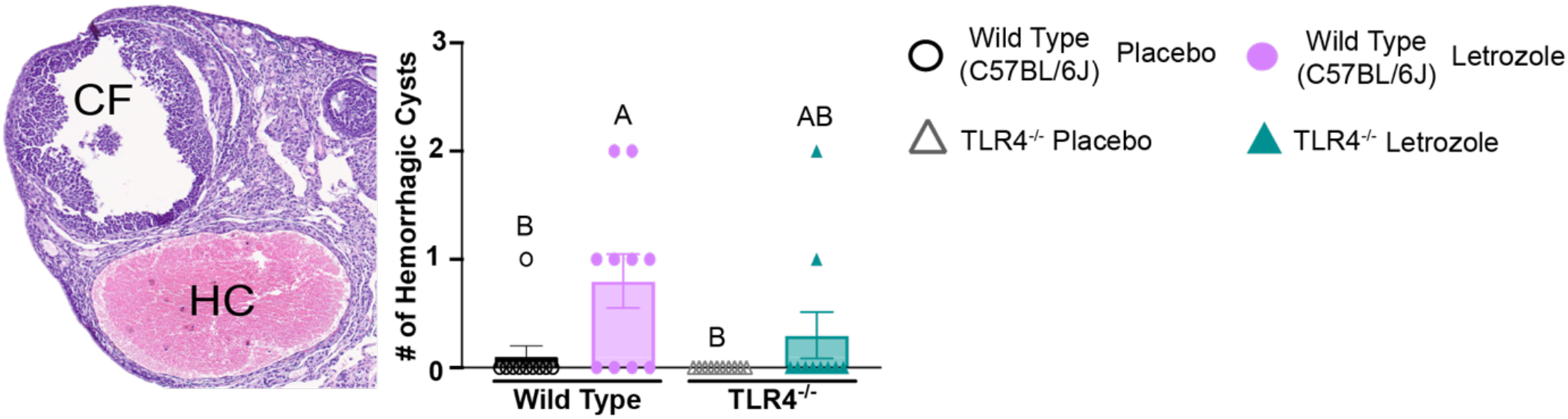
(Related to Figure 3) Ovarian histology. (A) Representative H&E-stained ovarian section (10 μm) from LET-treated WT mice showing hemorrhagic cyst (HC) morphological features. Data are presented as mean ± SEM and analyzed with two-way ANOVA followed by Tukey’s post-hoc test. Statistical significance was accepted at p < 0.05 and differences among groups are denoted by a connecting letter system, where groups sharing the same letter are not significantly different from each other, while groups with different letters are significantly different (p < 0.05).

## References

1. G. Bozdag, S. Mumusoglu, D. Zengin, E. Karabulut, B. O. Yildiz, The prevalence and phenotypic features of polycystic ovary syndrome: a systematic review and meta-analysis. Hum. Reprod. 31, 2841–2855 (2016).

2. R. Deswal, V. Narwal, A. Dang, C. S. Pundir, The Prevalence of Polycystic Ovary Syndrome: A Brief Systematic Review. J. Hum. Reprod. Sci. 13, 261 (2020).

3. S. Kulkarni, K. Gupta, P. Ratre, P. K. Mishra, Y. Singh, A. Biharee, S. Thareja, Polycystic ovary syndrome: Current scenario and future insights. Drug Discov. Today 28, 103821 (2023).

4. V. Unfer, E. Kandaraki, L. Pkhaladze, S. Roseff, M. H. Vazquez-Levin, A. S. Laganà, C. Shiao-Yng, M. I. M. Yap-Garcia, N. D. E. Greene, C. O. Soulage, A. Bevilacqua, S. Benvenga, D. Barbaro, B. Pintaudi, A. Wdowiak, C. Aragona, Z. Kamenov, M. Appetecchia, G. Porcaro, I. Hernandez Marin, F. Facchinetti, T. Chiu, O. Pustotina, O. Papalou, M. Nordio, T. Cantelmi, P. Cavalli, I. Vucenik, R. D’Anna, V. R. Unfer, S. Dinicola, S. Salehpour, A. Stringaro, M. Montaninno Oliva, M. Tugushev, N. Prapas, M. Bizzarri, M. S. B. Espinola, C. Di Lorenzo, A. C. Ozay, J. Nestler, When one size does not fit all: Reconsidering PCOS etiology, diagnosis, clinical subgroups, and subgroup-specific treatments. Endocr. Metab. Sci. 14, 100159 (2024).

5. H. F. Escobar-Morreale, Polycystic ovary syndrome: definition, aetiology, diagnosis and treatment. Nat. Rev. Endocrinol. 14, 270–284 (2018).

6. A. Acevedo-Rodriguez, A. S. Kauffman, B. D. Cherrington, C. S. Borges, T. A. Roepke, M. Laconi, Emerging insights into Hypothalamic-pituitary-gonadal (HPG) axis regulation and interaction with stress signaling. J. Neuroendocrinol. 30, e12590 (2018).

7. A. M. Moore, R. E. Campbell, The neuroendocrine genesis of polycystic ovary syndrome: A role for arcuate nucleus GABA neurons. J. Steroid Biochem. Mol. Biol. 160, 106–117 (2016).

8. A. S. Kauffman, V. G. Thackray, G. E. Ryan, K. P. Tolson, C. A. Glidewell-Kenney, S. J. Semaan, M. C. Poling, N. Iwata, K. M. Breen, A. J. Duleba, E. Stener-Victorin, S. Shimasaki, N. J. Webster, P. L. Mellon, A Novel Letrozole Model Recapitulates Both the Reproductive and Metabolic Phenotypes of Polycystic Ovary Syndrome in Female Mice1. Biol. Reprod. 93, 69, 1–12 (2015).

9. K. E. Kostroun, K. Goldrick, J. N. Mondshine, R. D. Robinson, E. Mankus, S. Reddy, Z. Wang, X. Song, J. F. Knudtson, Impact of updated international diagnostic criteria for the diagnosis of polycystic ovary syndrome. FS Rep. 4, 173–178 (2023).

10. S. Aboeldalyl, C. James, E. Seyam, E. M. Ibrahim, H. E.-D. Shawki, S. Amer, The Role of Chronic Inflammation in Polycystic Ovarian Syndrome-A Systematic Review and Meta-Analysis. Int. J. Mol. Sci. 22, 2734 (2021).

11. H. F. Escobar-Morreale, M. Luque-Ramírez, F. González, Circulating inflammatory markers in polycystic ovary syndrome: a systematic review and metaanalysis. Fertil. Steril. 95, 1048–1058.e1–2 (2011).

12. K. Azarbayjani, S. Jahanian Sadatmahalleh, A. Mottaghi, M. Nasiri, Association of dietary inflammatory index with C-reactive protein and interleukin-6 in women with and without polycystic ovarian syndrome. Sci. Rep. 14, 3972 (2024).

13. G. Pacheco-Sanchez, R. Herrera, M. D. Pisarska, R. Azziz, D. A. Nicholas, J. L. Chan, Multiracial Cytokine Profiling Reveals Immune Dysregulation not Chronic Inflammation in Polycystic Ovary Syndrome. J. Clin. Endocrinol. Metab., dgaf524 (2025).

14. A. Ascani, S. Torstensson, S. Risal, H. Lu, G. Eriksson, C. Li, S. Teschl, J. Menezes, K. Sandor, C. Ohlsson, C. I. Svensson, M. C. Karlsson, M. H. Stradner, B. Obermayer-Pietsch, E. Stener-Victorin, The role of B cells in immune cell activation in polycystic ovary syndrome. eLife 12, e86454 (2023).

15. C. Hu, B. Pang, Z. Ma, H. Yi, Immunophenotypic Profiles in Polycystic Ovary Syndrome. Mediators Inflamm. 2020, 5894768 (2020).

16. A. J. Duleba, A. Dokras, Is PCOS an inflammatory process? Fertil. Steril. 97, 7–12 (2012).

17. S. Mohammad, C. Thiemermann, Role of Metabolic Endotoxemia in Systemic Inflammation and Potential Interventions. Front. Immunol. 11, 594150 (2021).

18. Q. Zhu, H. Zhou, A. Zhang, R. Gao, S. Yang, C. Zhao, Y. Wang, J. Hu, R. Goswami, L. Gong, Q. Li, Serum LBP Is Associated with Insulin Resistance in Women with PCOS. PLOS ONE 11, e0145337 (2016).

19. B. Banaszewska, M. Siakowska, I. Chudzicka-Strugala, R. J. Chang, L. Pawelczyk, B. Zwozdziak, R. Spaczynski, A. J. Duleba, Elevation of markers of endotoxemia in women with polycystic ovary syndrome. Hum. Reprod. Oxf. Engl. 35, 2303–2311 (2020).

20. S. S. Ghosh, J. Wang, P. J. Yannie, S. Ghosh, Intestinal Barrier Dysfunction, LPS Translocation, and Disease Development. J. Endocr. Soc. 4, bvz039 (2020).

21. T. Kawai, M. Ikegawa, D. Ori, S. Akira, Decoding Toll-like receptors: Recent insights and perspectives in innate immunity. Immunity 57, 649–673 (2024).

22. Y. Wang, H. Xiao, Y. Liu, Q. Tong, Y. Yu, B. Qi, X. Bu, T. Pan, Y. Xing, Effects of Bu Shen Hua Zhuo formula on the LPS/TLR4 pathway and gut microbiota in rats with letrozole-induced polycystic ovary syndrome. Front. Endocrinol. 13 (2022).

23. Z.-P. Chang, G.-F. Deng, Y.-Y. Shao, D. Xu, Y.-N. Zhao, Y.-F. Sun, S.-Q. Zhang, R.-G. Hou, J.-J. Liu, Shaoyao-Gancao Decoction Ameliorates the Inflammation State in Polycystic Ovary Syndrome Rats via Remodeling Gut Microbiota and Suppressing the TLR4/NF-κB Pathway. Front. Pharmacol. 12, 670054 (2021).

24. B.-X. Gu, X. Wang, B.-L. Yin, H.-B. Guo, H.-L. Zhang, S.-D. Zhang, C.-L. Zhang, Abnormal expression of TLRs may play a role in lower embryo quality of women with polycystic ovary syndrome. Syst. Biol. Reprod. Med. 62, 353–358 (2016).

25. S. Akira, K. Takeda, Toll-like receptor signalling. Nat. Rev. Immunol. 4, 499–511 (2004).

26. R. Guo, Y. Zheng, J. Yang, N. Zheng, Association of TNF-alpha, IL-6 and IL-1beta gene polymorphisms with polycystic ovary syndrome: a meta-analysis. BMC Genet. 16, 5 (2015).

27. L. A. Velloso, F. Folli, M. J. Saad, TLR4 at the Crossroads of Nutrients, Gut Microbiota, and Metabolic Inflammation. Endocr. Rev. 36, 245–271 (2015).

28. P. D. Cani, J. Amar, M. A. Iglesias, M. Poggi, C. Knauf, D. Bastelica, A. M. Neyrinck, F. Fava, K. M. Tuohy, C. Chabo, A. Waget, E. Delmée, B. Cousin, T. Sulpice, B. Chamontin, J. Ferrières, J.-F. Tanti, G. R. Gibson, L. Casteilla, N. M. Delzenne, M. C. Alessi, R. Burcelin, Metabolic Endotoxemia Initiates Obesity and Insulin Resistance. Diabetes 56, 1761–1772 (2007).

29. L. Jia, C. R. Vianna, M. Fukuda, E. D. Berglund, C. Liu, C. Tao, K. Sun, T. Liu, M. J. Harper, C. E. Lee, S. Lee, P. E. Scherer, J. K. Elmquist, Hepatocyte Toll-like receptor 4 regulates obesity-induced inflammation and insulin resistance. Nat. Commun. 5, 3878 (2014).

30. Y. Zhao, G. Li, Y. Li, Y. Wang, Z. Liu, Knockdown of Tlr4 in the Arcuate Nucleus Improves Obesity Related Metabolic Disorders. Sci. Rep. 7, 7441 (2017).

31. D. V. Skarra, A. Hernández-Carretero, A. J. Rivera, A. R. Anvar, V. G. Thackray, Hyperandrogenemia Induced by Letrozole Treatment of Pubertal Female Mice Results in Hyperinsulinemia Prior to Weight Gain and Insulin Resistance. Endocrinology 158, 2988–3003 (2017).

32. S. Maurya, S. Tripathi, T. Arora, A. Singh, Adropin ameliorates reproductive dysfunctions in letrozole-induced PCOS mouse. Sci. Rep. 15, 8659 (2025).

33. P. Arroyo, B. S. Ho, L. Sau, S. T. Kelley, V. G. Thackray, Letrozole treatment of pubertal female mice results in activational effects on reproduction, metabolism and the gut microbiome. PLOS ONE 14, e0223274 (2019).

34. E. A. Coutinho, L. A. Esparza, J. Rodriguez, J. Yang, D. Schafer, A. S. Kauffman, Targeted inhibition of kisspeptin neurons reverses hyperandrogenemia and abnormal hyperactive LH secretion in a preclinical mouse model of polycystic ovary syndrome. Hum. Reprod. 39, 2089–2103 (2024).

35. L. A. Esparza, D. Schafer, B. S. Ho, V. G. Thackray, A. S. Kauffman, Hyperactive LH Pulses and Elevated Kisspeptin and NKB Gene Expression in the Arcuate Nucleus of a PCOS Mouse Model. Endocrinology 161, bqaa018 (2020).

36. A. Martin, S. Devkota, Hold the Door: Role of the Gut Barrier in Diabetes. Cell Metab. 27, 949–951 (2018).

37. C. A. Thaiss, M. Levy, I. Grosheva, D. Zheng, E. Soffer, E. Blacher, S. Braverman, A. C. Tengeler, O. Barak, M. Elazar, R. Ben-Zeev, D. Lehavi-Regev, M. N. Katz, M. Pevsner-Fischer, A. Gertler, Z. Halpern, A. Harmelin, S. Aamar, P. Serradas, A. Grosfeld, H. Shapiro, B. Geiger, E. Elinav, Hyperglycemia drives intestinal barrier dysfunction and risk for enteric infection. Science 359, 1376–1383 (2018).

38. C. Chelakkot, J. Ghim, S. H. Ryu, Mechanisms regulating intestinal barrier integrity and its pathological implications. Exp. Mol. Med. 50, 1–9 (2018).

39. J. J. Bunker, A. Bendelac, IgA responses to microbiota. Immunity 49, 211–224 (2018).

40. L. A. Zenewicz, IL-22 Binding Protein (IL-22BP) in the Regulation of IL-22 Biology. Front. Immunol. 12, 766586 (2021).

41. A. Fantou, E. Lagrue, T. Laurent, L. Delbos, S. Blandin, A. Jarry, G. Beriou, C. Braudeau, N. Salabert, E. Marin, A. Moreau, J. Podevin, A. Bourreille, R. Josien, J. C. Martin, IL-22BP production is heterogeneously distributed in Crohn’s disease. Front. Immunol. 13 (2022).

42. H. Shi, M. V. Kokoeva, K. Inouye, I. Tzameli, H. Yin, J. S. Flier, TLR4 links innate immunity and fatty acid–induced insulin resistance. J. Clin. Invest. 116, 3015–3025 (2006).

43. D. Pal, S. Dasgupta, R. Kundu, S. Maitra, G. Das, S. Mukhopadhyay, S. Ray, S. S. Majumdar, S. Bhattacharya, Fetuin-A acts as an endogenous ligand of TLR4 to promote lipid-induced insulin resistance. Nat. Med. 18, 1279–1285 (2012).

44. L. Jia, C. R. Vianna, M. Fukuda, E. D. Berglund, C. Liu, C. Tao, K. Sun, T. Liu, M. J. Harper, C. E. Lee, S. Lee, P. E. Scherer, J. K. Elmquist, Hepatocyte Toll-like receptor 4 regulates obesity-induced inflammation and insulin resistance. Nat. Commun. 5, 3878 (2014).

45. K.-A. Kim, W. Gu, I.-A. Lee, E.-H. Joh, D.-H. Kim, High Fat Diet-Induced Gut Microbiota Exacerbates Inflammation and Obesity in Mice via the TLR4 Signaling Pathway. PLOS ONE 7, e47713 (2012).

46. M. Saberi, N.-B. Woods, C. de Luca, S. Schenk, J. C. Lu, G. Bandyopadhyay, I. M. Verma, J. M. Olefsky, Hematopoietic Cell-Specific Deletion of Toll-like Receptor 4 Ameliorates Hepatic and Adipose Tissue Insulin Resistance in High-Fat-Fed Mice. Cell Metab. 10, 419–429 (2009).

47. P. F. Svendsen, L. Nilas, S. Madsbad, J. J. Holst, Incretin hormone secretion in women with polycystic ovary syndrome: roles of obesity, insulin sensitivity, and treatment with metformin. Metabolism. 58, 586–593 (2009).

48. K. Sridharan, G. Sivaramakrishnan, Expanding therapeutic horizons: glucagon-like peptide-1 receptor agonists and sodium glucose transporter-2 inhibitors in poly cystic ovarian syndrome: a comprehensive review including systematic review and network meta-analysis of randomized clinical trials. Diabetol. Metab. Syndr. 17, 168 (2025).

49. I. Gunesli, E. Ulug, A. A. Pinar, O. Portakal, B. O. Yildiz, Fasting and postprandial oxytocin and incretin dynamics in women with polycystic ovary syndrome and healthy controls. Eur. J. Endocrinol. 193, 255–261 (2025).

50. Y. Luan, L. Zhang, Y. Peng, Y. Li, R. Liu, C. Yin, Immune regulation in polycystic ovary syndrome. Clin. Chim. Acta 531, 265–272 (2022).

51. L. Geng, X. Yang, J. Sun, X. Ran, D. Zhou, M. Ye, L. Wen, R. Wang, M. Chen, Gut Microbiota Modulation by Inulin Improves Metabolism and Ovarian Function in Polycystic Ovary Syndrome. Adv. Sci. 12, 2412558 (2025).

52. R. Wu, S. Fujii, N. K. Ryan, K. H. Van der Hoek, M. J. Jasper, I. Sini, S. A. Robertson, R. L. Robker, R. J. Norman, Ovarian leukocyte distribution and cytokine/chemokine mRNA expression in follicular fluid cells in women with polycystic ovary syndrome. Hum. Reprod. 22, 527–535 (2007).

53. H. Zhang, X. Wang, J. Xu, Y. Zhu, X. Chen, Y. Hu, IL-18 and IL-18 binding protein concentration in ovarian follicular fluid of women with unexplained infertility to PCOS during in vitro fertilization. J. Reprod. Immunol. 138, 103083 (2020).

54. Y. Liu, H. Liu, Z. Li, H. Fan, X. Yan, X. Liu, J. Xuan, D. Feng, X. Wei, The Release of Peripheral Immune Inflammatory Cytokines Promote an Inflammatory Cascade in PCOS Patients via Altering the Follicular Microenvironment. Front. Immunol. 12, 685724 (2021).

55. S. Torstensson, A. Ascani, S. Risal, H. Lu, A. Zhao, A. Espinosa, E. Lindgren, M. H. Johansson, G. Eriksson, M. Barakat, M. C. I. Karlsson, C. Svensson, A. Benrick, E. Stener-Victorin, Androgens Modulate the Immune Profile in a Mouse Model of Polycystic Ovary Syndrome. Adv. Sci. Weinh. Baden-Wurtt. Ger. 11, e2401772 (2024).

56. H.-Y. Guan, H.-X. Xia, X.-Y. Chen, L. Wang, Z.-J. Tang, W. Zhang, Toll-Like Receptor 4 Inhibits Estradiol Secretion via NF-κB Signaling in Human Granulosa Cells. Front. Endocrinol. 12, 629554 (2021).

57. Q. Yang, Q. Wan, Z. Wang, Curcumin mitigates polycystic ovary syndrome in mice by suppressing TLR4/MyD88/NF-κB signaling pathway activation and reducing intestinal mucosal permeability. Sci. Rep. 14, 29848 (2024).

58. D. Zhang, L. Zhang, F. Yue, Y. Zheng, R. Russell, Serum zonulin is elevated in women with polycystic ovary syndrome and correlates with insulin resistance and severity of anovulation. Eur. J. Endocrinol. 172, 29–36 (2015).

59. A. Zak-Gołąb, P. Kocełak, M. Aptekorz, M. Zientara, L. Juszczyk, G. Martirosian, J. Chudek, M. Olszanecka-Glinianowicz, Gut microbiota, microinflammation, metabolic profile, and zonulin concentration in obese and normal weight subjects. Int. J. Endocrinol. 2013, 674106 (2013).

60. J. Wang, S. S. Ghosh, S. Ghosh, Curcumin improves intestinal barrier function: modulation of intracellular signaling, and organization of tight junctions. Am. J. Physiol. - Cell Physiol. 312, C438–C445 (2017).

61. L. Lindheim, M. Bashir, J. Münzker, C. Trummer, V. Zachhuber, B. Leber, A. Horvath, T. R. Pieber, G. Gorkiewicz, V. Stadlbauer, B. Obermayer-Pietsch, Alterations in Gut Microbiome Composition and Barrier Function Are Associated with Reproductive and Metabolic Defects in Women with Polycystic Ovary Syndrome (PCOS): A Pilot Study. PLOS ONE 12, e0168390 (2017).

62. K. Tremellen, K. Pearce, Dysbiosis of Gut Microbiota (DOGMA) – A novel theory for the development of Polycystic Ovarian Syndrome. Med. Hypotheses 79, 104–112 (2012).

63. X. Qi, C. Yun, L. Sun, J. Xia, Q. Wu, Y. Wang, L. Wang, Y. Zhang, X. Liang, L. Wang, F. J. Gonzalez, D. Patterson, H. Liu, L. Mu, Z. Zhou, Y. Zhao, R. Li, P. Liu, C. Zhong, Y. Pang, C. Jiang, J. Qiao, Gut microbiota–bile acid–interleukin-22 axis orchestrates polycystic ovary syndrome. Nat. Med. 25, 1225–1233 (2019).

64. J. Kempski, A. D. Giannou, K. Riecken, L. Zhao, B. Steglich, J. Lücke, L. Garcia-Perez, K.-F. Karstens, A. Wöstemeier, M. Nawrocki, P. Pelczar, M. Witkowski, S. Nilsson, L. Konczalla, A. M. Shiri, J. Kempska, R. Wahib, L. Brockmann, P. Huber, A.-C. Gnirck, J.-E. Turner, D. E. Zazara, P. C. Arck, A. Stein, R. Simon, A. Daubmann, J. Meiners, D. Perez, T. Strowig, P. Koni, A. A. Kruglov, G. Sauter, J. R. Izbicki, A. H. Guse, T. Rösch, A. W. Lohse, R. A. Flavell, N. Gagliani, S. Huber, IL22BP Mediates the Antitumor Effects of Lymphotoxin Against Colorectal Tumors in Mice and Humans. Gastroenterology 159, 1417–1430.e3 (2020).

65. A. M. Hammer, N. L. Morris, A. R. Cannon, O. M. Khan, R. C. Gagnon, N. V. Movtchan, I. van Langeveld, X. Li, B. Gao, M. A. Choudhry, Interleukin-22 Prevents Microbial Dysbiosis and Promotes Intestinal Barrier Regeneration Following Acute Injury. Shock Augusta Ga 48, 657–665 (2017).

66. J. Kempski, A. D. Giannou, K. Riecken, L. Zhao, B. Steglich, J. Lücke, L. Garcia-Perez, K.-F. Karstens, A. Wöstemeier, M. Nawrocki, P. Pelczar, M. Witkowski, S. Nilsson, L. Konczalla, A. M. Shiri, J. Kempska, R. Wahib, L. Brockmann, P. Huber, A.-C. Gnirck, J.-E. Turner, D. E. Zazara, P. C. Arck, A. Stein, R. Simon, A. Daubmann, J. Meiners, D. Perez, T. Strowig, P. Koni, A. A. Kruglov, G. Sauter, J. R. Izbicki, A. H. Guse, T. Rösch, A. W. Lohse, R. A. Flavell, N. Gagliani, S. Huber, IL22BP Mediates the Antitumor Effects of Lymphotoxin Against Colorectal Tumors in Mice and Humans. Gastroenterology 159, 1417–1430.e3 (2020).

67. P. J. Torres, D. V. Skarra, B. S. Ho, L. Sau, A. R. Anvar, S. T. Kelley, V. G. Thackray, Letrozole treatment of adult female mice results in a similar reproductive phenotype but distinct changes in metabolism and the gut microbiome compared to pubertal mice. BMC Microbiol. 19, 57 (2019).

68. S. T. Kelley, D. V. Skarra, A. J. Rivera, V. G. Thackray, The Gut Microbiome Is Altered in a Letrozole-Induced Mouse Model of Polycystic Ovary Syndrome. PLOS ONE 11, e0146509 (2016).

69. S. Gupta, S. L. Gupta, A. Singh, N. Oswal, V. Bal, S. Rath, A. George, S. Basu, IgA Determines Bacterial Composition in the Gut. Crohns Colitis 360 5, otad030 (2023).

70. Y. Yang, J. Cheng, C. Liu, X. Zhang, N. Ma, Z. Zhou, W. Lu, C. Wu, Gut microbiota in women with polycystic ovary syndrome: an individual based analysis of publicly available data. eClinicalMedicine 77 (2024).

71. L. Wang, J. Zhou, H.-J. Gober, W. T. Leung, Z. Huang, X. Pan, C. Li, N. Zhang, L. Wang, Alterations in the intestinal microbiome associated with PCOS affect the clinical phenotype. Biomed. Pharmacother. Biomedecine Pharmacother. 133, 110958 (2021).

72. Y. Li, Y. Zhu, D. Li, W. Liu, Y. Zhang, W. Liu, C. Zhang, T. Tao, Depletion of gut microbiota influents glucose metabolism and hyperandrogenism traits of mice with PCOS induced by letrozole. Front. Endocrinol. 14 (2023).

73. H. Luck, S. Khan, J. H. Kim, J. K. Copeland, X. S. Revelo, S. Tsai, M. Chakraborty, K. Cheng, Y. Tao Chan, M. K. Nøhr, X. Clemente-Casares, M.-C. Perry, M. Ghazarian, H. Lei, Y.-H. Lin, B. Coburn, Okrainec, T. Jackson, S. Poutanen, H. Gaisano, J. P. Allard, D. S. Guttman, M. E. Conner, S. Winer, D. A. Winer, Gut-associated IgA+ immune cells regulate obesity-related insulin resistance. Nat. Commun. 10, 3650 (2019).

74. A. Khajouei, E. Hosseini, T. Abdizadeh, M. Kian, S. Ghasemi, Beneficial effects of minocycline on the ovary of polycystic ovary syndrome mouse model: Molecular docking analysis and evaluation of TNF-α, TNFR2, TLR-4 gene expression. J. Reprod. Immunol. 144, 103289 (2021).

75. A. Otoo, A. Czika, J. Lamptey, J.-P. Yang, Q. Feng, M.-J. Wang, Y.-X. Wang, Y.-B. Ding, Emodin improves glucose metabolism and ovarian function in PCOS mice via the HMGB1/TLR4/NF-κB molecular pathway. Reprod. Camb. Engl. 166, 323–336 (2023).

76. S. Wold, M. Sjöström, L. Eriksson, PLS-regression: a basic tool of chemometrics. Chemom. Intell. Lab. Syst. 58, 109–130 (2001).

77. R. P. Simmons, E. P. Scully, E. E. Groden, K. F. Benedict, J. J. Chang, K. Lane, J. Lifson, E. Rosenberg, D. A. Lauffenburger, M. Altfeld, HIV-1 infection induces strong production of IP-10 through TLR7/9-dependent pathways. AIDS Lond. Engl. 27, 2505–2517 (2013).

78. D. A. Nicholas, E. A. Proctor, M. Agrawal, A. C. Belkina, S. C. Van Nostrand, L. Panneerseelan-Bharath, A. R. Jones, F. Raval, B. C. Ip, M. Zhu, J. Cacicedo, C. Habib, N. Sainz-Rueda, L. Persky, P. G. Sullivan, B. E. Corkey, C. M. Apovian, P. A. Kern, D. A. Lauffenburger, B. S. Nikolajczyk, Fatty Acid Metabolites Combine with Reduced β Oxidation to activate Th17 Inflammation in Human Type 2 Diabetes. Cell Metab. 30, 447–461.e5 (2019).

79. I. Barroeta-Espar, L. D. Weinstock, B. G. Perez-Nievas, A. C. Meltzer, M. Siao Tick Chong, A. C. Amaral, M. E. Murray, K. L. Moulder, J. C. Morris, N. J. Cairns, J. E. Parisi, V. J. Lowe, R. C. Petersen, J. Kofler, M. D. Ikonomovic, O. López, W. E. Klunk, R. P. Mayeux, M. P. Frosch, L. B. Wood, T. Gomez-Isla, Distinct cytokine profiles in human brains resilient to Alzheimer’s pathology. Neurobiol. Dis. 121, 327–337 (2019).

80. L. B. Wood, A. R. Winslow, E. A. Proctor, D. McGuone, D. A. Mordes, M. P. Frosch, B. T. Hyman, D. Lauffenburger, K. M. Haigis, Identification of neurotoxic cytokines by profiling Alzheimer’s disease tissues and neuron culture viability screening. Sci. Rep. 5, 16622 (2015).

81. G. P. Sanchez, M. Lopez, L. M. Velez, I. Tamburini, N. Ujagar, J. A. Angulo, G. De Robles, H. Choi, J. Arriola, R. Kapadia, A. B. Zonderman, M. K. Evans, C. Jang, M. M. Seldin, D. A. Nicholas, Comparative Analysis of White and African American Groups Reveals Unique Lipid and Inflammatory Features of Diabetes. J. Racial Ethn. Health Disparities, doi: 10.1007/s40615-025-02642-z (2025).

